# SieveAI: Development of an Automated extensible and customisable drug discovery pipeline and its validation

**DOI:** 10.1101/2025.04.20.648820

**Authors:** Vishal Kumar Sahu, Anshuman Sand, Sangeeta Ballav, Vajrini Raman, Shuchi Nagar, Amit Ranjan, Soumya Basu

**Affiliations:** Cancer and Translational Research Centre, Dr. D. Y. Patil Biotechnology and Bioinformatics Institute, Dr. D. Y. Patil Vidyapeeth, Pune-411033, India; Bioinformatics Centre, Dr. D. Y. Patil Biotechnology and Bioinformatics Institute, Dr. D. Y. Patil Vidyapeeth, Pune-411033, Maharashtra, India

**Author notes:** Corresponding Authors: **Dr. Soumya Basu**, Associate Professor, Cancer and Translational Research Centre, Dr. D. Y. Patil Biotechnology and Bioinformatics Institute, Dr. D. Y. Patil Vidyapeeth, Tathawade, Pune - 411033, India, **Dr. Amit Ranjan**, Associate Professor, Cancer and Translational Research Centre, Dr. D. Y. Patil Biotechnology and Bioinformatics Institute, Dr. D. Y. Patil Vidyapeeth, Tathawade, Pune - 411033, India.

**Keywords:** Molecular docking automation, Protein-ligand docking, Protein-RNA docking, RNA-ligand docking

## Abstract

Systematic Interaction Evaluation and Virtual Enhancement Analysis Interface (SieveAI) is an automated drug discovery pipeline developed to enhance the efficiency of virtual screening and computer-aided drug discovery processes. The molecular docking workflow encompasses acquiring, modeling, and pre-processing of molecular structure files, conducting docking with various algorithms, and subsequent analysis and interpretation of the outcomes by visualising or tabulating the results. While several open-source software tools are available to assist these operations at different steps of molecular docking, they often necessitate manual user intervention at every stage. To streamline and automate this extensive manual process and develop a comprehensive solution, we have developed an innovative, fully extensible, molecular docking pipeline SieveAI (©L-129927/2023). The same has been demonstrated in this manuscript. SieveAI works with a range of open-source libraries, packages, and programs to facilitate automated drug discovery using established programs and software. The package is accessible at https://miRNA.in/SieveAI.

## Introduction

The rapid development and evolution of computational power have significantly transformed the landscape of life sciences research particularly drug discovery. This progress has enabled researchers to evaluate the continuously growing number of molecular structures in the last few decades. Additionally, the emergence of AI-based modeling approaches holds great promise in addressing challenges related to structure elucidation, marking a significant advancement in the field. As a testament to this progress, the Research Collaboratory for Structural Bioinformatics (RCSB) repository, as of October, 2024, boasts a staggering collection of 225,946 three dimensional structures of RNA, DNA, and protein [1]. Impressively, this repository witnesses an annual growth of approximately 9% in structure submissions [2]. Further, advancing algorithms like AlphaFold and ESMFold have predicted over 600 million protein structures providing a breakthrough in the long standing problem of protein folding with enhanced accuracy [3–6]. On the other side, there is availability of millions of drug-like molecules and compounds belonging to categories of small molecules, natural products, and synthetic peptides having therapeutic potential [7,8].

In traditional drug discovery, the identification of novel therapeutic compounds involves a laborious and time-consuming process requiring extensive experimental testing and validation. However, with the advent of automated pipelines, researchers can leverage computational methods, high-throughput screening techniques, and machine learning algorithms to expedite the discovery of potential drug candidates. One of the main components of drug discovery involves molecular interaction analysis that deals with the interactions among receptors and ligands. These structures undergo numerous optimisation, preparatory, interaction analysis, and interpretation steps using complex computational algorithms which are tedious, time consuming, and error prone with a lot of manual intervention [9,10].

Automated technologies, such as computational algorithms, and high-throughput screening, now streamline and accelerate various stages of the drug discovery pipeline and they dramatically enhance efficiency, speed, and precision in developing novel therapeutics. These advancements reduce time and costs while increasing scalability, allowing rapid screening of thousands of compounds, thus improving the robustness and reproducibility of experimental results. Consequently, automation holds great promise for addressing unmet medical needs and expediting the delivery of innovative treatments [9].

Our proposed drug discovery pipeline, Systematic Interaction Evaluation and Virtual Enhancement Analysis Interface (SieveAI), represents a paradigm shift in the realm of drug discovery. SieveAI pipeline offers a cutting-edge platform that leverages computational algorithms to expedite virtual drug screening against proteins, RNAs like miRNA, lncRNAs etc, lipids, DNA. As the demand for novel therapeutics continues to escalate in response to emerging diseases and to fulfill medical needs, the pipeline is imperative to streamline and accelerate the drug discovery process. SieveAI extensively upgrade the landscape of pharmaceutical research aiming to streamline and accelerate the process of drug discovery and development by reducing cost, time, and identification of drug-target related toxicity. The source code of SieveAI can be accessed at https://www.miRNA.in/SieveAI.

## 2 Methodology

### 2.1 Components and Architecture of the SieveAI Pipeline

The SieveAI pipeline is designed with a modular architecture that includes a core package and customizable plugins. The core package contains essential classes and processes for structure processors or managers and docking processors or managers while the plugins integrating external software or programs like AutoDockVINA, ChimeraX, HDockLite, PatchDock etc allow for customization at different stages of the pipeline. SieveAI integrates functionalities from various open-source programs and algorithms, providing a seamless and automated workflow for virtual drug screening. Further it relies on the open source packages like biopython to effectively process and analyse biomolecules.

### 2.2 Input and Retrieval of Molecular Structures

SieveAI can accept molecular structures manually or retrieved automatically from public databases through identifiers listed as prescribed in documentation. For manual input, researchers can curate structures such as RNA, DNA, proteins, or chemical compounds from databases like PDB, PubChem, KEGG Compounds, Coconut, or UniProt. The core module of SieveAI can be customized to retrieve protein structures from UniProt or select the best available PDB structure from AlphaFold if necessary [4,11].

### 2.3 Preprocessing and Docking Preparation

Some of the docking algorithms like AutoDock VINA require preprocessing by converting the input PDB file format to PDBQT format, calculation of binding grid, and input parameter file as basic requirement to start molecular docking of macromolecule with small molecules [12].

### 2.4 Docking Algorithms and Execution

While these steps are redundant in other packages analysing macromolecule to macromolecule interactions like PatchDock, HADDock, or HDockLite and it can take PDB structures directly as input to perform protein-protein, protein-RNA, protein-DNA, RNA-RNA, or DNA-RNA based molecular interaction analysis [13–15]. Basically, there are two types of docking approaches, blind docking where the whole receptor molecule is checked for a possible binding site of the provided ligand and site-specific docking where a specific site or potential binding pocket is provided to the algorithms to perform the docking. The implementation of the approach depends on the docking algorithm to perform site specific docking or blind docking; the input can be customized by updating relevant plugins within the SieveAI during the docking preparation step.

SieveAI uses plugins to perform docking analysis using different algorithms, for example AutoDock VINA, HDock, ChimeraX, VMD, and OpenBabel. It provides input parameters like program specific parameters, input file names, or output file names to these docking algorithms, and performs docking in parallel using the multiprocessing module of python [16].

### 2.5 Rescoring and Reanalysis

Since, most of the docking algorithms and packages are optimised for protein-protein docking there becomes further requirement for rescoring involving traditional or machine learning based algorithms. One of the cases being, performing molecular docking of RNA and small molecule using AutoDock VINA that requires rescoring to optimize the molecule docking interaction scores [17–19]. SieveAI provides a plugin to implement rescoring using AnnapuRNA to rescore RNA-molecule docking. More rescoring methods can be added to the pipeline through the plugin customisation approach.

### 2.6 Post-docking Interaction Analysis

The molecular docking analysis is further subjected to analysis using a post processing plugin for analysis involving ChimeraX and VMD-Python to calculate Hydrogen bonds, polar and non-polar solvent accessible surface area, contacts, and salt-bridges [20–22]. These parameters in addition to scores provided by the docking programs are further ranked using composite index method.

### 2.7 Ranking and Validation

The ranks for interactions were calculated by assigning rank based weights to the score, number of contacts, and hydrogen bonds of the predicted docking conformations. These weights were further refined with the composite index construction method [23–25]. In the context of protein-RNA interaction analysis, composite index construction involves integrating multiple variables like scores, number of contacts, hydrogen bonds, solvent accessible surface area (SASA) of the receptor fragment, SASA of ligand fragment, and distance between the geometric centers of the fragments obtained from docking analysis into a single metric for ranking the conformations. The advantage of using a composite index is that it allows for a more comprehensive evaluation of interactions compared to relying on individual criteria alone. It takes into account the relative importance of each factor. This approach helps prioritize and identify the most significant protein-RNA interacting conformers. SieveAI pipeline streamlines manual input, file handling, and tabulation therefore the validation of SieveAI working was performed by comparing pipeline output with manually processed outcomes.

## 3 Results and Discussion

### 3.1 Architecture and Functionality of SieveAI

SieveAI was developed as an automated drug discovery pipeline to streamline the molecular docking process. The architecture of SieveAI is modular, consisting of a core package and several customizable plugins. The core package includes essential classes and processes that handle configuration, preprocessing, docking execution, and post-processing analysis. The plugin-based architecture allows for customization at various stages of the pipeline, enabling researchers to integrate specific tools and algorithms suited to their needs.

SieveAI integrates functionalities from several open-source programs and algorithms, including AutoDock VINA, HDockLite, ChimeraX, VMD, and OpenBabel. This integration allows SieveAI to perform a wide range of molecular docking tasks, from protein-ligand docking to more complex interactions like protein-RNA and RNA-ligand docking. The pipeline’s design emphasizes flexibility and extensibility, allowing users to add new plugins and extend the core functionalities as needed.

SieveAI code base comprises core modules dealing with processes and validation of molecules with manual code or open-source libraries summarised in Figure 1. The core consists of configuration to deal with the user input from web interface, command line interface, or through extended codebase. Further, these configurations are passed on to the preprocessor blocks to preprocess molecules like trimming water molecules, natural ligands, or additional structure in the crystal structure. Further these structures are passed to the plugin where the conversion of file formats and preparation of parameter files is handled by the algorithm specific plugin followed by molecule docking interaction. After the processing and docking the processing blocks send it for the rescoring if configured else results from the docking program are parsed and interaction analysis is performed leading to docking pose ranking. These ranks are further tabulated and provided as docking result format. Optionally these results can be assessed by LLM based analysis for intricate input from AI.

**Figure 1:**
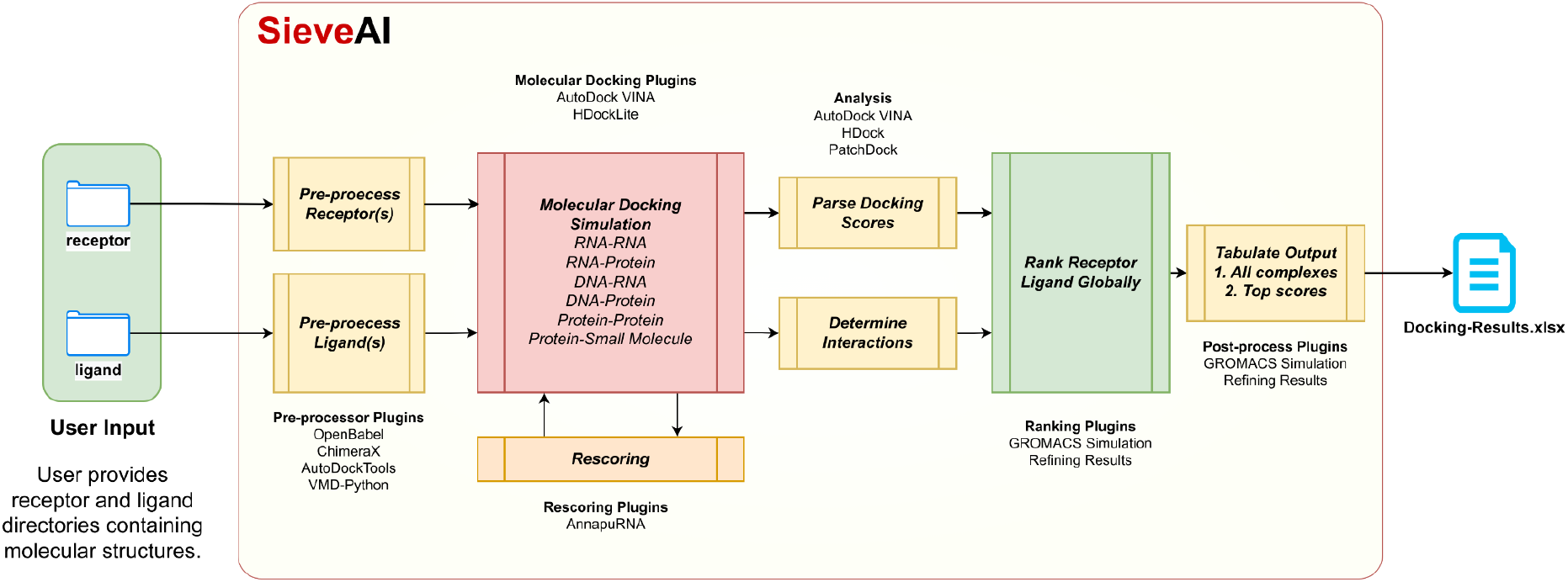
Workflow involved in SieveAI to automate molecular docking

SieveAI can be extended to traditional or modern artificial intelligence based algorithms for docking, web based docking programs, and other programs that are not covered here.

SieveAI pipeline is compatible with all operating systems that support execution of python programming and the software and packages required in the pipeline to carry out molecular docking and analysis. SieveAI can be hosted on a server to centrally perform molecular interaction simulation or analysis.

### 3.2 Code Organisation and Extendibility

**Figure 1:**
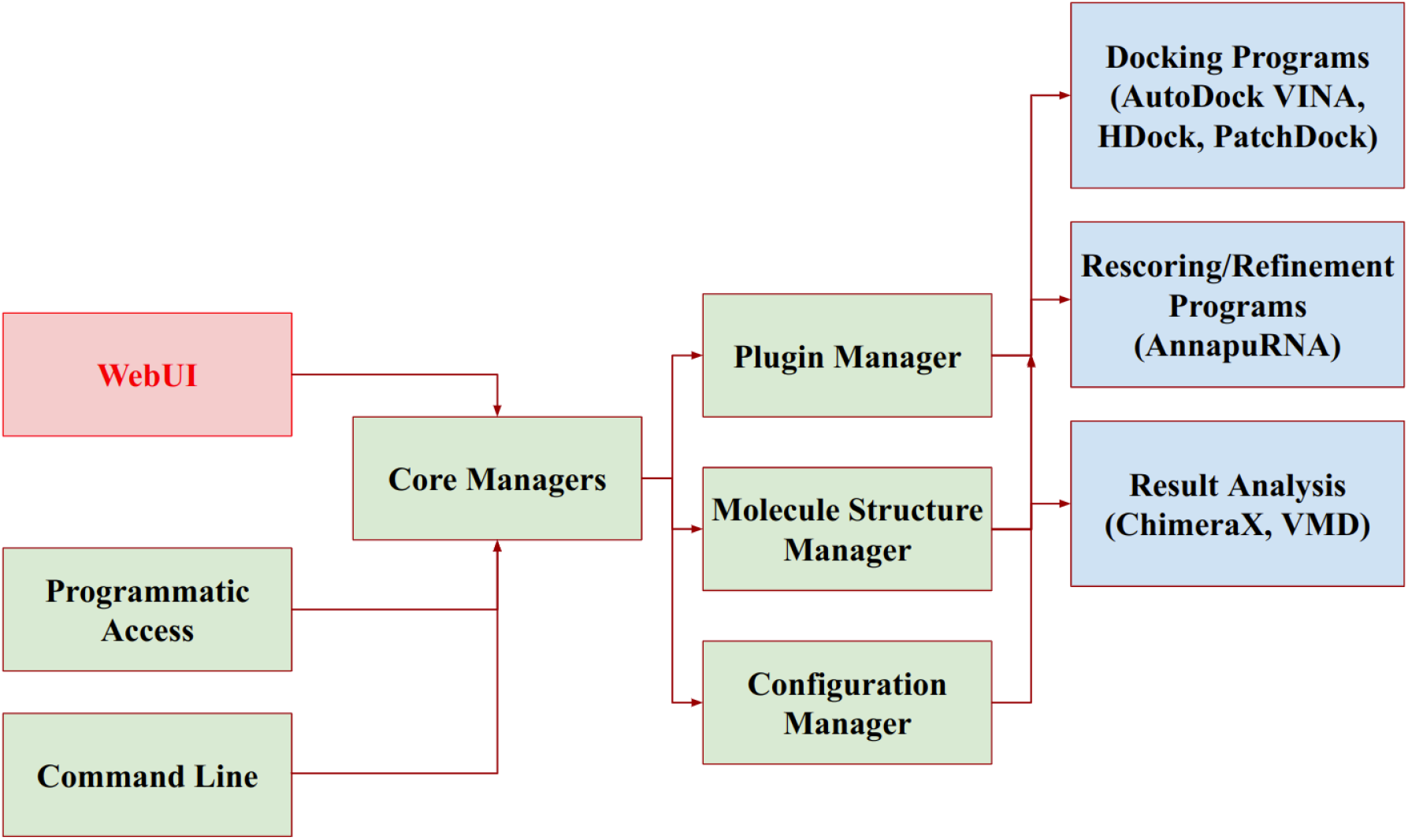
Plugin Interactions and management of input and output. WebUI and WebServer are made available through different programs in a decoupled manner.

**Table 1:** List of existing plugins in SieveAI architecture to connect with and control external program.

| Name of the plugin | Purpose | External Program Controlled | Reference |
| --- | --- | --- | --- |
| Vina | Molecular docking of protein-ligand, RNA-ligand, lipid-ligand, and DNA-ligands; Grid preparation | AutoDock VINA 1.2.3 | [12] |
| ChimeraX | Interaction Analysis, Publication Oriented Graphic Generation | ChimeraX | [20] |
| VMDPy | Interaction analysis and other calculations | VMD Python | [21] |
| HDockLite | Docking of macromolecules like protein-protein, protein-NA, or NA-NA (NA: Nucleic acids like RNA or DNA) | HDock | [14] |
| Converter | Conversion of molecules from one | OpenBabel, | [27,28] |
|  | file format to other file format | MGLTools,<br>AutoDock Tools |  |
| AnnapuRNA | Rescoring of RNA-ligand docking results from AutoDock VINA | AnnapuRNA | [17] |
| StructureSync | Retrieval of molecular structures from public databases using given file input | Manual Scripts to Retrieve Molecule Structures from Pubchem and RCSB PDB |  |

SieveAI can retrieve molecular structures from the given list and additionally integrate pathway information like KEGG or WikiPathways from where it can extract gene information and later retrieve the protein structures with the help of UniProt by converting gene ids into PDB ids and downloading best structure in terms of completeness and availability.

### 3.3 Efficiency calculation of SieveAI Pipeline

To validate the efficiency and accuracy of the SieveAI pipeline, we compared the outcomes of the automated process with manually conducted docking processes. The validation involved performing molecular docking of 10 complexes manually and using SieveAI. The manual docking process took an average of 3 minutes per complex, whereas SieveAI completed the docking in one minute per complex on the same system, without utilizing parallel computing or multiprocessing capabilities.

The results from SieveAI were found to be consistent with those obtained manually, demonstrating the accuracy and reliability of the pipeline. The automated process significantly reduced the time required for docking and minimized the potential for human error. This validation underscores the effectiveness of SieveAI in streamlining the molecular docking workflow and enhancing overall efficiency.

SieveAI pipeline maintains input parameters through configuration with default input parameters to maintain the reproducibility. It saves custom parameters when input parameters for any program or software are customized. The results and analysis are subject to change with the change in the input parameters.

In case of molecular docking using AutoDock VINA a random seed value is provided to the VINA docking program as input parameter that results in approximately the similar results. Docking interaction prediction is totally a random process and results in random conformers.

The pipeline substitutes the manual repeating steps involved in virtual screening or molecule drug discovery, therefore the validation results were manually compared by carrying out docking of few conformers and comparing the outcomes of SieveAI pipeline.

Manual docking of 10 complexes took an average of 3 minutes per complex while when it was done using SieveAI took one complex per minute on the same system without involving parallel computing or multiprocessing.

### 3.4 Composite Index Ranking (CIR) Approach and Result Output of SieveAI

The ranking approach used in the SieveAI helps in normalizing the ranking of parameters or outcomes of different properties of interactions like score, H-Bonds, contacts, and salt-bridges that are different in terms of their units and values. Therefore the implementation of composite index based ranking provides an added advantage to consider all the aspects before ranking the conformers.

Score, H-bonds, contacts, and other parameters are tabulated after their calculation and output is preserved in a raw format and provided in excel format for visual interpretation. Further the ranking output is also provided to manually analyse the results.

**Figure 1:**
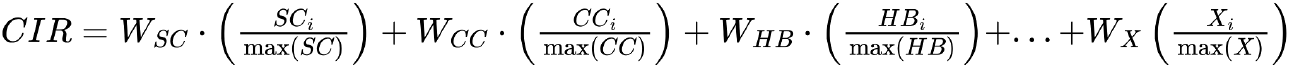
Formula for composite index ranking (CIR) used by SieveAI for ranking docking results. (i = i^th^ complex; CC Contacts Count; HB Hydrogen Bond Counts, X Other matrices like SASA of receptor and ligand available)

### 3.5 Validation of SieveAI Pipeline

#### 3.5.1 In silico Validation: Molecular Docking of Protein p53 with FDA-approved drugs

We performed molecular docking of TP53 protein (PDB ID 1TUP) with 1,947 FDA-approved molecules to demonstrate the effectiveness and efficiency of SieveAI. The timing was monitored to obtain outcome in a system equipped with Intel Core i5 8th Gen, 8 GB RAM, and 512 GB SSD (Table 2 and Supplementary File 2). In comparison to manual and other docking pipelines, SieveAI calculates Contacts, Hydrogen Bonds and additionally ranks them based on Composite Index ranking method providing a complete workflow for molecular docking approach.

**Table 2:** Molecular docking analysis of TP53 (PDB ID 1TUP) with DrugBank drugs.

| **Compo**<br>**site**<br>**Rank** | **VINA**<br>**Score** | **Drug**<br>**Molecule** | **Total**<br>**HBonds** | **Total**<br>**Contacts** | **HBond Residues** |
| --- | --- | --- | --- | --- | --- |
| 1 | -10.0967 | Diosmin<br>(DB08995) | 6 | 126 | SER:96,ARG:196,ARG:196,ASN:200,UNL:1,UNL:1 |
| 2 | -8.9254 | Hesperidin<br>(DB04703) | 5 | 116 | SER:96,VAL:172,ARG:196,ARG:196,UNL:1 |
| 3 | -8.6674 | Indinavir<br>(DB00224) | 5 | 138 | THR:140,ARG:196,ARG:196,GLY:199,UNL:1 |
| 4 | -9.1019 | Glipizide<br>(DB01067) | 5 | 105 | ARG:267,ARG:267,THR:231,UNL:1,UNL:1 |
| 5 | -9.3593 | Lifitegrast<br>(DB11611) | 4 | 121 | THR:140,ARG:196,GLY:199,ASN:200 |
| 6 | -8.8918 | Gliquidone<br>(DB01251) | 4 | 115 | SER:99,THR:231,UNL:1,UNL:1 |
| 7 | -8.418 | NADH<br>(DB00157) | 5 | 109 | LYS:139,ARG:196,UNL:1,UNL:1,UNL:1 |
| 8 | -8.8989 | Glyburide<br>(DB01016) | 4 | 108 | SER:99,THR:231,UNL:1,UNL:1 |
| 9 | -8.1992 | Doxorubicin<br>(DB00997) | 5 | 120 | HIS:233,HIS:233,UNL:<br>1,UNL:1,UNL:1 |
| 10 | -9.8963 | Diacetyl<br>benzoyl<br>lathyrol<br>(DB11260) | 4 | 97 | SER:96,ARG:196,ARG<br>:196,GLY:199 |

**Table 3:** Statistics of Outcomes from SieveAI.

|  | SieveAI Pipeline | Manual Docking |
| --- | --- | --- |
| Average Time Taken for Single Docking Using AutoDock VINA | 1 Min | 1 Min |
| Average Time Taken By AutoDock VINA for complete analysis till ranking | 3 Min | > 10 Min |
| Parallel Docking Analysis | Yes | No |
| Manual Error | No | Yes |

#### 3.5.2 In silico Validation: Molecular Docking of hsa-miR-9-5p and FDA Approved drugs from DrugBank using SieveAI

To demonstrate the flexibility and extensibility of the SieveAI pipeline we performed molecular docking of human miR-9-5p and 1,597 FDA-approved drugs (small molecules) obtained from DrugBank. This work also helped us to narrow down repurposed drugs in TNBC. The docking was performed using Vina plugin that uses AutoDock VINA 1.2.3 for molecular docking, followed by rescoring using ML based algorithm AnnapuRNA to rescore RNA-ligand docking, and analyzed using ChimeraX. The ranked docking conformers of drugs were then subjected to literature search and few of the drugs were reported to interact with miR-9-5p in breast cancer. Mitomycin (DB00305) and colchicine (DB01394) are representative drugs that we found amongst the drugs that interact with hsa-miR-9-5p (Table and Supplementary File 3) with top ranking VINA Score of −7.4608 kcal/mol and AnnapuRNA Score of −415.505 kcal/mol with Amikacin and predicted formation of 10 hydrogen bonds and 110 contacts.

**Table 4:** Molecular docking and rescore results of top 10 drugs for docking with hsa-miR-9-5p.

| Com<br>posit<br>e<br>Rank | VINA<br>Score | AnnapuR<br>NA Score | Drug Molecule | Tot<br>al<br>HB<br>on<br>ds | Total<br>Cont<br>acts | HBond Residues |
| --- | --- | --- | --- | --- | --- | --- |
| 1 | -7.4608 | -415.505 | Amikacin<br>(DB00479) | 10 | 110 | U:4,G:15,UNL:1,UNL:1,<br>UNL:1,UNL:1,UNL:1,U<br>NL:1,UNL:1,UNL:1 |
| 2 | -7.9162 | -395.832 | Acarbose<br>(DB00284) | 9 | 105 | U:4,A:14,A:14,C:16,UN<br>L:1,UNL:1,UNL:1,UNL:<br>1,UNL:1 |
| 3 | -8.7308 | -400.019 | Paromomycin<br>(DB01421) | 8 | 115 | U:4,C:16,UNL:1,UNL:1,<br>UNL:1,UNL:1,UNL:1,U<br>NL:1 |
| 4 | -8.4992 | -368.38 | Framycetin<br>(DB00452) | 7 | 97 | U:4,G:7,C:16,UNL:1,UN<br>L:1,UNL:1,UNL:1 |
| 5 | -8.5995 | -291.732 | Ribostamycin<br>(DB03615) | 8 | 98 | U:4,U:4,C:16,UNL:1,UN<br>L:1,UNL:1,UNL:1,UNL:<br>1 |
| 6 | -7.7799 | -318.608 | Plazomicin<br>(DB12615) | 7 | 95 | U:4,G:7,C:16,UNL:1,UN<br>L:1,UNL:1,UNL:1 |
| 7 | -7.9457 | -320.554 | Micronomicin<br>(DB13274) | 6 | 134 | G:7,A:10,C:16,UNL:1,U<br>NL:1,UNL:1 |
| 8 | -7.9729 | -311.853 | Kanamycin<br>(DB01172) | 9 | 81 | G:7,G:7,A:14,G:15,C:16,<br>UNL:1,UNL:1,UNL:1,U<br>NL:1 |
| 9 | -7.9501 | -291.021 | Dibekacin<br>(DB13270) | 6 | 112 | G:6,G:7,C:16,UNL:1,UN<br>L:1,UNL:1 |
| 9 | -8.0887 | -321.995 | Pentosan<br>polysulfate<br>(DB00686) | 10 | 80 | U:4,G:7,A:14,A:14,A:14,<br>G:15,C:16,C:16,UNL:1,<br>UNL:1 |

### 3.7 Advantages of SieveAI Pipeline

| Pack<br>age<br>Name | Int<br>egr<br>ate<br>d<br>We<br>b<br>UI | Inte<br>grat<br>ed<br>Web<br>Serv<br>er | Protei<br>n<br>Down<br>loading | Liga<br>nd<br>Retri<br>eval | Mole<br>cule<br>Prep<br>arati<br>on | Integrati<br>on of<br>Multiple<br>Docking<br>Program<br>s | Docking<br>Analysis | Type<br>of<br>Docki<br>ng | Rank<br>ing<br>Appro<br>ach | Par<br>allel<br>ism | Cust<br>omiz<br>abilit<br>y | Wor<br>kflo<br>w |
| --- | --- | --- | --- | --- | --- | --- | --- | --- | --- | --- | --- | --- |
| Sieve<br>AI | No | Yes | Autom<br>atic<br>from<br>PDB<br>and<br>UniPro<br>t | Requ<br>ired<br>and<br>Auto<br>mate<br>d | Auto<br>Dock<br>VINA<br>,<br>HDoc<br>kLite | Automate<br>d using<br>VMD and<br>ChimeraX | Protein-Ligand,<br>Protein-Protein,<br>Protein-RNA,<br>Protein-DNA,<br>RNA-Li<br>gand,<br>DNA-Li<br>gand | Comp<br>osite<br>Index | Yes<br>for<br>large-<br>scale<br>docki<br>ng | Para<br>llel<br>dock<br>ing<br>appro<br>ach | Open<br>Sour<br>ce<br>Scrip<br>t | 3<br>work<br>flows<br>are<br>avail<br>able |
| Ca<br>ver<br>Do<br>ck | Yes | No | PDB | Yes | Chim<br>era | Supports<br>AutoDock<br>Vina | Focused<br>on<br>channels<br>and<br>tunnels<br>in<br>proteins | Protei<br>n-Lig<br>and | Path-<br>based<br>scorin<br>g | Limi<br>ted | Low | 1<br>prima<br>ry<br>work<br>flow |
| mz<br>Do<br>ck | No | Yes | From<br>various<br>sources | Limit<br>ed | PyM<br>OL<br>integr<br>ation | No | Limited | Protei<br>n-Lig<br>and,<br>Small<br>molec<br>ule | Basic<br>energ<br>y-bas<br>ed | Limi<br>ted | Medi<br>um | Custo<br>miza<br>ble<br>with<br>plugi<br>ns |
| Gli<br>de | Yes | Yes | PDB | Schr<br>odin<br>ger<br>datab<br>ases | Maest<br>ro | No | Protein-Ligand | Glide<br>Score | Medi<br>um | Prop<br>rieta<br>ry | 1<br>prim<br>ary<br>work<br>flow | No |
| Au | No | No | Manual | Man | Auto | Yes | Protein- | Energy | No | High | Fully | No |
| to<br>Do<br>ck<br>To<br>ols |  |  |  | ual | Dock |  | Ligand | y-bas<br>ed |  | (Scri<br>ptin<br>g-ba<br>sed) | custo<br>miza<br>ble |  |

#### Limitations and Future Prospects

SieveAI shows efficiency by reducing manual intervention and produces reliability by controlling the programs attested in the pipeline with given user input and avails extensibility in terms of incorporating new workflow and involvement of new software even AI driven. However, the current version of SieveAI has its limitations as it utilises external libraries and programs for molecular docking. The workflow doesn’t consider docking of metalloproteins while protein preparation therefore requires integration of other programs.

The pipeline currently lacks built-in support for docking metalloproteins, which are essential in many biological processes. Metalloproteins involve metal ions that require specialized handling in molecular docking because standard force fields and docking tools may not appropriately capture metal coordination and binding effects. Additionally it requires further modification for site-specific docking and ranked results modification as it totally depends on external libraries and programs.

While the pipeline offers a plugin for rescoring docking results including RNA-ligand, Protein-ligand, RNA-RNA, or Protein-Protein, rescoring remains statistically dependent on score, contacts, and number of hydrogen bonds. Apart from these, WebUI or GUI may considerably make it easy to use.

## Conclusion

The development of the Systematic Interaction Evaluation and Virtual Enhancement Analysis Interface (SieveAI) marks a significant advancement to aid in the field of drug discovery. Our findings demonstrate that SieveAI can efficiently automate the complex and repetitive processes involved in molecular docking. It can assist from molecular structure retrieval to interaction analysis and ranking providing an end-to-end solution. SieveAI reduces manual intervention, thereby minimizing errors and significantly accelerates the drug discovery pipeline by leveraging computational algorithms and integrating multiple open-source tools including modern machine learning based algorithms. The results indicate that SieveAI can handle various types of molecular interactions, including protein-ligand, protein-RNA, RNA-ligand, or nucleic acid-nucleic acid based interactions with a higher reproducibility and fidelity. SieveAI’s approach involving pre-processing, docking, rescoring, result curation, and ranking showcases its potential to streamline drug discovery workflow. Additionally, real-world applications of SieveAI can be explored through collaborations with pharmaceutical companies and research institutions. By continuously updating and refining the pipeline, SieveAI can play a pivotal role in addressing emerging medical needs and accelerating the development of innovative therapeutics.

## Supplementary Files

Supplementary File 1: FDA Approved Drugs used in the study with their Identifiers and descriptions

Supplementary File 2: SieveAI Assisted Molecular Docking of hsa-miR-9-5p with FDA Approved Drugs

Supplementary File 3: SieveAI Assisted Molecular Docking of p53 with FDA Approved Drugs

## Acknowledgements

Authors acknowledge Dr. D. Y. Patil Biotechnology and Bioinformatics Institute, Pune, India for research infrastructure and the Department of Science & Technology (DST), Government of India for providing DST-FIST grant (SR/FST/LS-I/2017/70) to support research infrastructure and instruments. VKS acknowledges the Council of Scientific and Industrial Research (CSIR), New Delhi, India for CSIR-SRF fellowship (File No.: 09/1340(11487)/2021-EMR-I). SB acknowledges intramural grant of Dr. D. Y. Patil Vidyapeeth (DPU), Pune, India (DPU/644-43/2021).

